# Competitions for tyrosine breakdown: In synthetic microbial communities and between a gut microbial pathway and a human pathway

**DOI:** 10.1101/2025.07.07.663581

**Authors:** Madison Mitchem, Clint W. Pauling, Dhara D. Shah

**Affiliations:** Biodesign Center for fundamental and applied microbiomics, Arizona State University, Tempe, AZ 85281, USA; School of Life Sciences, Arizona State University, Tempe, AZ 85281, USA; School of Mathematical and Natural Sciences, Arizona State University, Glendale, AZ 85306, USA

## Abstract

Tyrosine, a versatile amino acid that undergoes diverse transformations to produce both beneficial and detrimental metabolites. The modulation of these metabolites results from direct competition among different metabolic pathways responsible for the breakdown of tyrosine whether it be the competition between distinct microbes or the rivalry between a microbe and its host. The fight between microbes for the available tyrosine might drive potential changes to the communities present in various environments. In contrast, if the similar contest for tyrosine is presented between a gut microbial pathway and a human pathway, it can hold potential to affect the human health. In this work, we present various metabolic outcomes of tyrosine within synthetic microbial communities which are prominently driven by specific enzyme activities of tyrosine breakdown pathways. Additionally, we developed a metabolic diversion of the human tyrosine breakdown pathway facilitated by a gut microbial enzyme. This approach holds promise as a novel strategy to develop potential therapeutic interventions in future for addressing metabolic disorders like tyrosinemias (I, II, III), hawkinsinuria, and alkaptonuria, associated with the human tyrosine breakdown pathway.

## INTRODUCTION

Tyrosine, is a proteinogenic and a versatile amino acid which can be metabolized either oxidatively^1^ or reductively^2^. In aerobic organisms, oxidative tyrosine catabolism involves a five step pathway that provides fumarate and acetoacetate (Fig. 1), which can ultimately enter into the TCA cycle and can be utilized as an energy source^1^. In contrast, the main reductive pathway for tyrosine breakdown that generally occur in anaerobic organisms, starts with the conversion of L-tyrosine to 4-hydroxyphenyl pyruvate (HPP) that ultimately gets converted to 4-hydroxypropionate^2^. However, those are not the only routes of tyrosine breakdown or metabolism. Tyrosine metabolism can produce numerous metabolites with a great variation and diversity^3,4^. Some of these metabolites are crucial for the organisms that produce them. For instance, homogentisate (HG) is necessary for photosynthetic organisms, and tyrosine-based neurotransmitters like dopamine and tyramine are necessary for humans. Whereas other metabolites may be detrimental or harmful to the organism that produces them or to other nearby organisms. Additionally, in the context of organisms which are part of the human gut microbiome, such molecules produced by the members of the community in the gut can affect the health of the host^5–7^. When a variety of tyrosine breakdown pathways are present within the same environment, the abundances of metabolites can be driven by the direct competition for substrate between enzymes of these metabolic pathways. It is crucial to understand such crosstalk within different microbial communities due to the multitudes of reactions that generate numerous tyrosine metabolites. Investigations like this can possibly help understand the metabolic outputs of microbial communities that are not solely based on gene annotations, but also on the functional efficiencies or fluxes of these metabolic pathways which can rely significantly on enzymes of the pathways. Here, we tackle this question by looking at select enzymes involved in tyrosine metabolism from three separate environmental microbial community members. Additionally, we utilized a gut microbial enzyme to bifurcate the tyrosine catabolism pathway that is similar to the human tyrosine breakdown pathway. The human tyrosine breakdown pathway involves five steps catalyzed by five different enzymes^8^. Due to mutations or abnormal activities in one or more enzymes involved in this tyrosine breakdown pathway, accumulation of harmful intermediates or products can occur, which can lead to five different types of metabolic disorders. These are; type I, II and III tyrosinemias, hawkinsinuria, and alkaptonuria^1,9–18^. To help patients with such metabolic disorders, the pathway needs to be inhibited. Currently, there is an FDA approved therapy available for the type I tyrosinemia, which is the most morbid of all known disorders, that occur due to errors within tyrosine breakdown^9,10,12,18–20^. This therapy includes the administration of the drug nitisinone (NTBC) which inhibits the enzyme 4-hydroxyphenylpyruvate dioxygenase (HPPD) that catalyzes the second step in the tyrosine catabolic pathway (Fig. 1). It is the first committed step of this pathway because the first reaction catalyzed by tyrosine aminotransferase (TAT) is reversible.

**Fig. 1:**
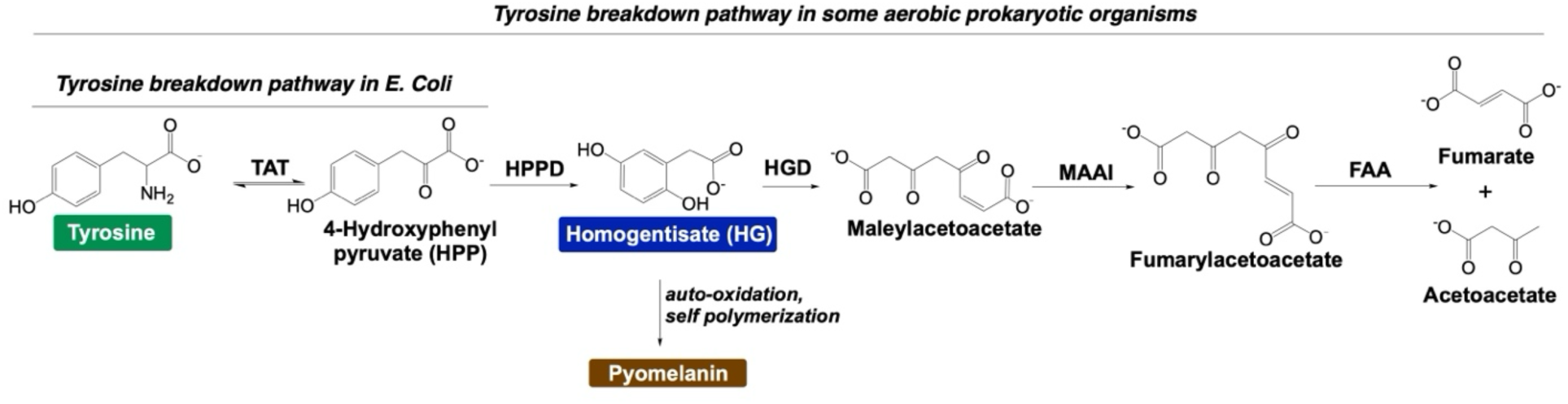
Tyrosine catabolism pathway from aerobic organisms. In the pathway, tyrosine is converted to fumarate and acetoacetate with five successive steps.

This inhibition, although used primarily to treat type I tyrosinemia, has the ability to help patients with three out of five abnormal tyrosine breakdown disorders mentioned above^1,9,10,12,18–20^. However, due the reversibility of the first step, the inhibition increases blood tyrosine concentrations that can result into many physiological abnormalities, e.g., visual impairments and IQ suppression over time^21–24^. This increase in blood tyrosine concentrations can be troublesome because it is an extremely insoluble molecule. To this end, we hypothesized creating a bifurcation in the human tyrosine metabolism pathway that can possibly lead to the breakdown of tyrosine without the production of harmful metabolites of the homogentisate dependent tyrosine breakdown pathway (Fig. 1) and could allow the constant utilization of tyrosine and hence may obstruct the increase in blood tyrosine levels. Utilizing synthetic communities created with *E. coli* strains and by simulating pathways of tyrosine breakdown that are found in gut microbes and in humans, we show the evidence of pathway diversions. Our results show that these metabolic rerouting arise due to, 1) the direct competition between enzymes from the members of the environmental microbial communities, and 2) the competition between a gut microbial enzyme and a human enzyme. Additionally, rather than blocking the tyrosine pathway that can increase insoluble tyrosine in the blood, utilizing a gut microbial enzyme to divert the harmful pathway and creating such continuous clearance system for tyrosine from the blood might help with developing a better therapeutic strategy in future.

## RESULTS AND DISCUSSION

### Pyomelanin serves as the indicator for secreted homogentisate

To test if a metabolic diversion could be achieved in microbial communities, we employed a phenotypic assay. An *E. coli* strain harboring the enzyme HPPD is able to produce a brown pigment when grown in LB media ^20,25,26^. *E. coli* is an excellent assay system here because it only has the first enzyme of the tyrosine breakdown pathway, known as tyrosine aminotransferase (TAT) that coverts tyrosine to 4-hydroxyphenylpyruvate (4-HPP) (Fig. 1). In addition, *E. coli* cells are able to transport tyrosine and its metabolites across the cell^25,26^. Here, we utilized an *E. coli* strain harboring an enzyme 4-hydroxyphenylpyruvate dioxygenase (HPPD) (Fig. 2). HPPD catalyzes the second step of the tyrosine catabolism pathway (Fig. 1). When this strain is given tyrosine, it first gets converted to 4-HPP and then to homogentisate (HG) due to the HPPD that is provided within the plasmid. Homogentisate can be secreted outside the cell, where it gets oxidized and self-polymerized to produce a brown pigment called pyomelanin^25,26^. This pigment can be measured at 405 nm. We tested *E. coli* BL21(DE3) for the pigment production due to the expression of the enzyme 4-hydroxypheylpyruvate dioxygenase (HPPD) from an environmental microbe (Fig. 2).

**Fig. 2:**
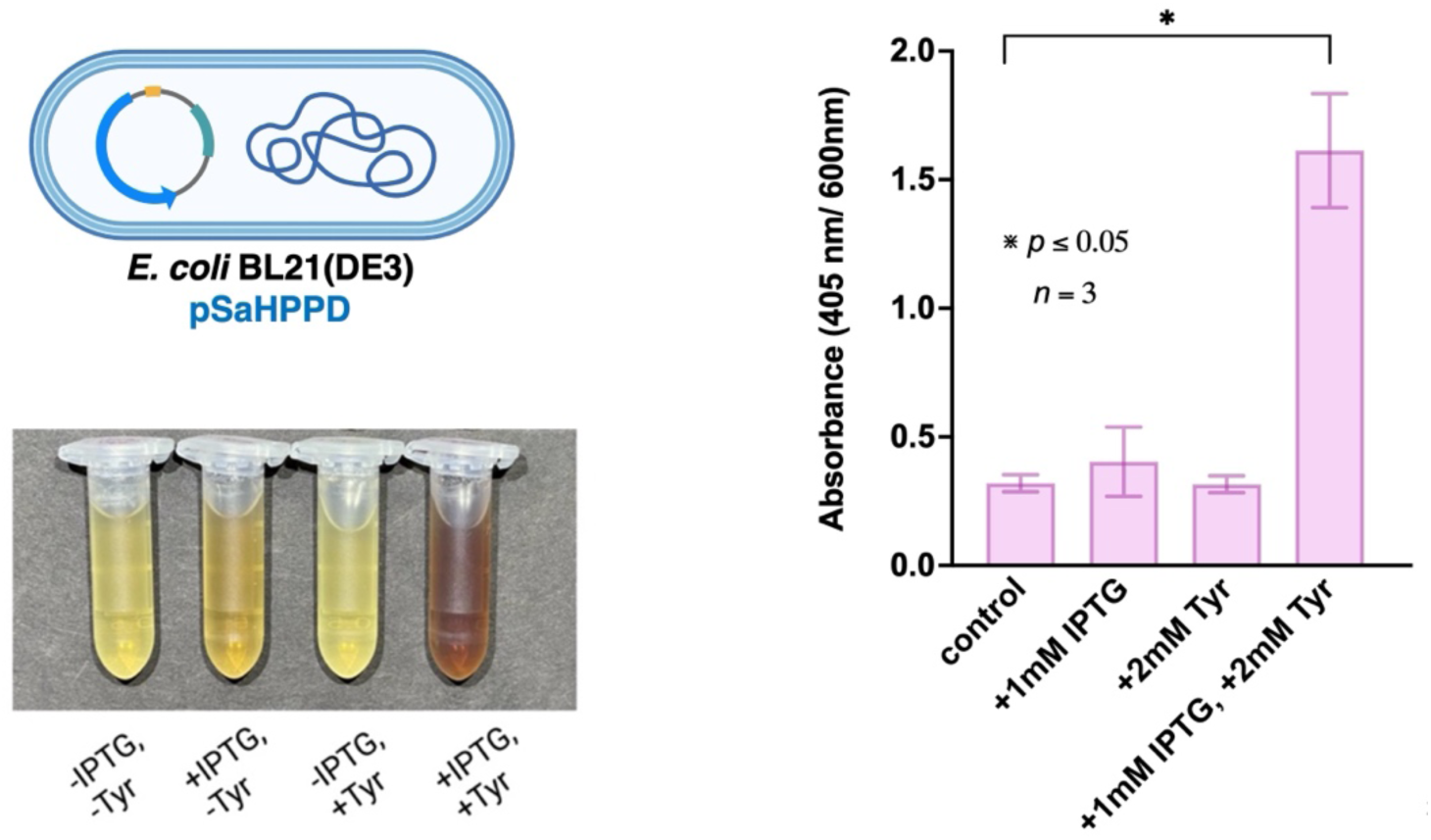
Pyomelanin serves as the indicator for secreted homogentisate. Pigment production in cultures of *E. coli* BL21(DE3) harboring SaHPPD under four different conditions can be seen. Quantification of pyomelanin production was carried out by normalizing pyomelanin absorbance by cell density for each strain in four different conditions. ANOVA test was used to calculate statistical significance and p values, n=3.

Figure 2 depicts pigment production due the presence of HPPD in *E. coli* BL21(DE3) cells. We observed the most pigment when these cultures were induced and had an additional source of the substrate tyrosine (Fig. 2, +IPTG, +Tyr). There was some pigment production when cultures were induced without the addition of the substrate, and we believe this is due to the additional minimal source of tyrosine found in the rich media (LB) used (Fig. 2, +IPTG). Figure 2 graph represents the quantification of the pigmentation in above mentioned cultures. Although, we didn’t see any difference in the growth rates of these cultures, we normalized the absorbance coming from the pigment via OD_600_ to eliminate any small differences in the number of cells that produce the pigment. There is a significant amount of pigment production in cultures that received both tyrosine and IPTG, specially for BL21(DE3) strain (Fig. 2 graph).

### Pigmentation is proportional to ratios of *E. coli* strains harboring different enzymes for tyrosine metabolism

To understand the allocation of tyrosine as a resource in complex microbial communities, we started with two *E. coli* strains differing only by the tyrosine metabolizing enzyme present on a plasmid. For this, we chose 4-hydroxyphenylpyruvate dioxygenase (SaHPPD) from *Streptomyces avermitilis*^27–29^ and hydroxymandelate synthase (AoHMS) from *Amycolatopsis orientalis*^30,31^ (Fig. 3 and Fig. 4). Both *S. avermitilis* and *A. orientalis* are environmental microbes and have the ability to produce a variety of natural products. For example, *S. avermitilis* produces avermectins which are utilized as antiparaitic, insecticide and antihelminitic drugs^32–34^. Whereas vancomycin, a peptide antibiotic was first isolated from *A. orientalis*^35^. A not proteinogenic amino acid 4-hydroxyphenylglycine (pHPG) is needed for the production of vancomycin and hydroxymandelate (HMA), the product of the reaction catalyzed by AoHMS is the precursor for pHPG^36^. In contrast, *S. avermitilis* produces multiple pigments like melanin that are produced from tyrosine^37^. To understand the competition in utilizing tyrosine within the community of such environmental microbes, we created a synthetic community of *E. coli* strains harboring and differing only in the presence of tyrosine metabolism enzymes (Fig. 3 and Fig. 4).

**Fig. 3:**
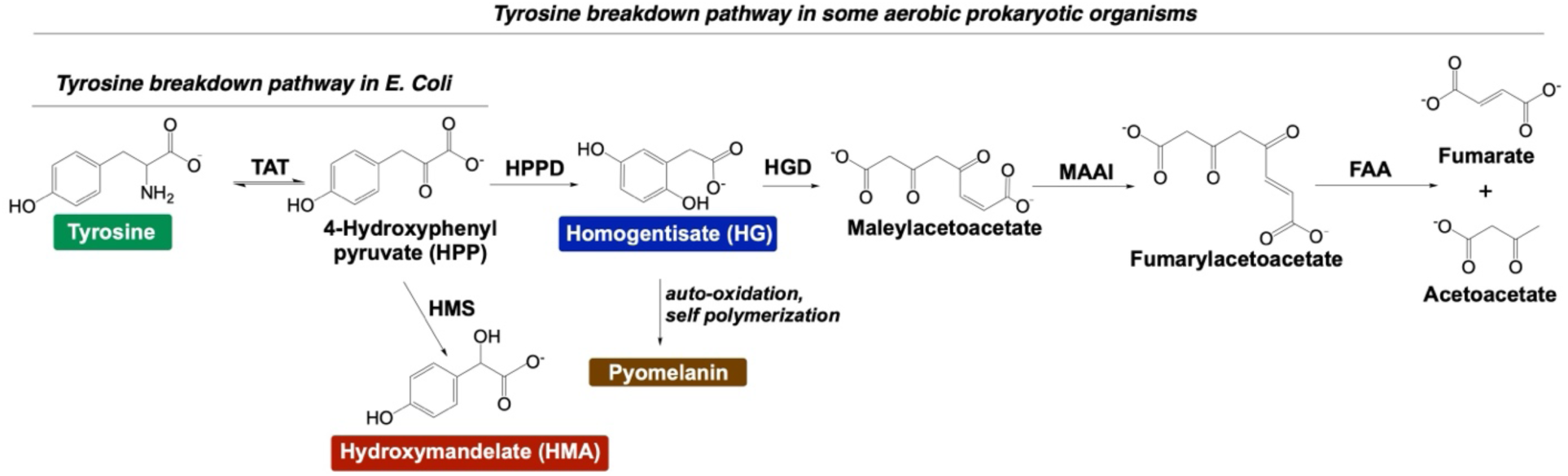
Tyrosine catabolism pathway diversion with one enzyme. The figure depicts the possible route for redirecting the tyrosine breakdown pathway by adding one additional enzyme involved in tyrosine metabolism. Here, hydroxymandelate synthase (HMS) can convert tyrosine to hydroxymandelate (HMA).

**Fig. 4:**
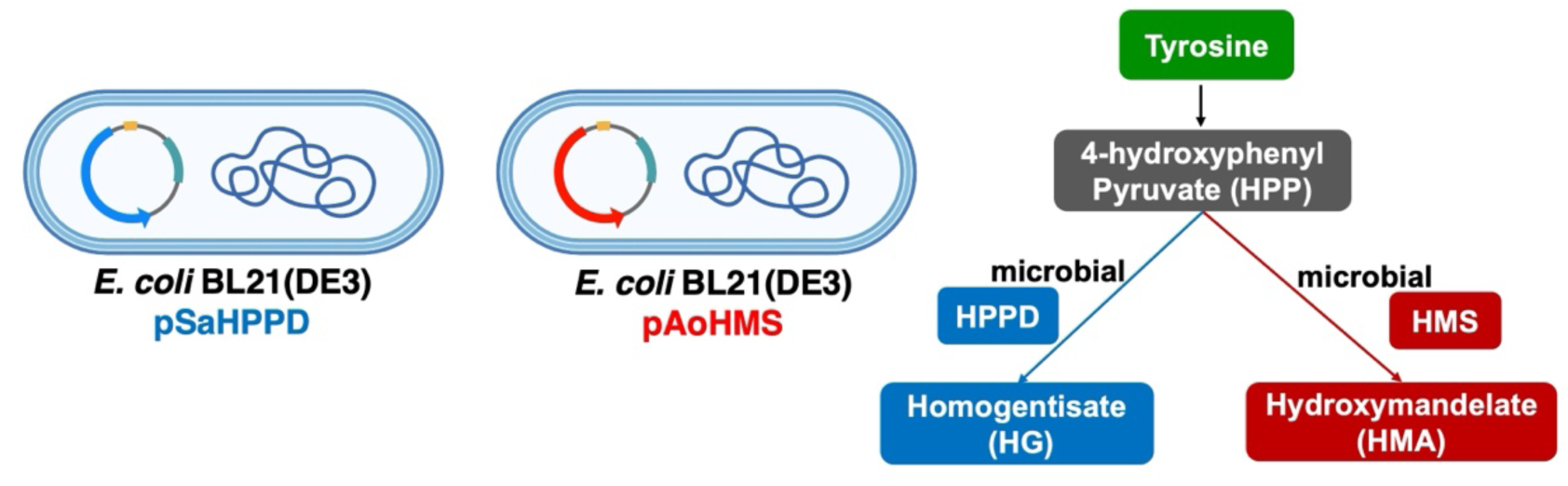
Generation of *E. coli* strains for Tyrosine catabolism pathway diversion with one enzyme. The figure depicts *E. coli* strains designed for redirecting the tyrosine breakdown pathway by adding one additional enzyme involved in tyrosine metabolism. Here, hydroxymandelate synthase (HMS) can convert tyrosine to hydroxymandelate (HMA).

The competition between SaHPPD and AoHMS proceeds via an intermediate 4-HPP (Fig. 3 and Fig. 4). When *E. coli* BL21(DE3) cells harboring either SaHPPD or AoHMS were cultivated individually or as mixtures of different ratios, the competition between pathways was visible through the change in the pyomelanin production (Fig. 5). Figure 5 shows the pigmentation in spent media of various cultures of *E. coli* BL21(DE3) cells harboring either SaHPPD or AoHMS or mixtures of the two at various ratios. The product of AoHMS is HMA that does not produce pyomelanin hence there was no pigmentation in the AoHMS culture. In contrast, homogentisate (HG), the product of SaHPPD, can produce pyomelanin hence a dark brown pigmentation was seen in the spent media of SaHPPD culture. Interestingly, when the cultures of AoHMS and SaHPPD were mixed at various ratios and then allowed to grow with induction and addition of tyrosine, the resulting cultures showed pigmentation levels that correlated very well with the initial ratios of two mixed *E. coli* strains, where higher amount of added *E. coli* cells with SaHPPD showed higher pigmentation (Fig. 5). As expected, the highest pigmentation is seen with SaHPPD culture followed by 4 SaHPPD:1 AoHMS, 3 SaHPPD:2 AoHMS, 2 SaHPPD:3 AoHMS, and then 1 SaHPPD:4 AoHMS. Culture ratio of 1 SaHPPD:1 AoHMS, showed pigmentation that was between the pigmentation level of 3 SaHPPD:2 AoHMS and 2 SaHPPD:3 AoHMS. This result showed that the direct competition between two enzymatic activities was being translated in the pigmentation differences which are ultimately driven due to changes in the metabolic output of this synthetic community.

**Fig. 5:**
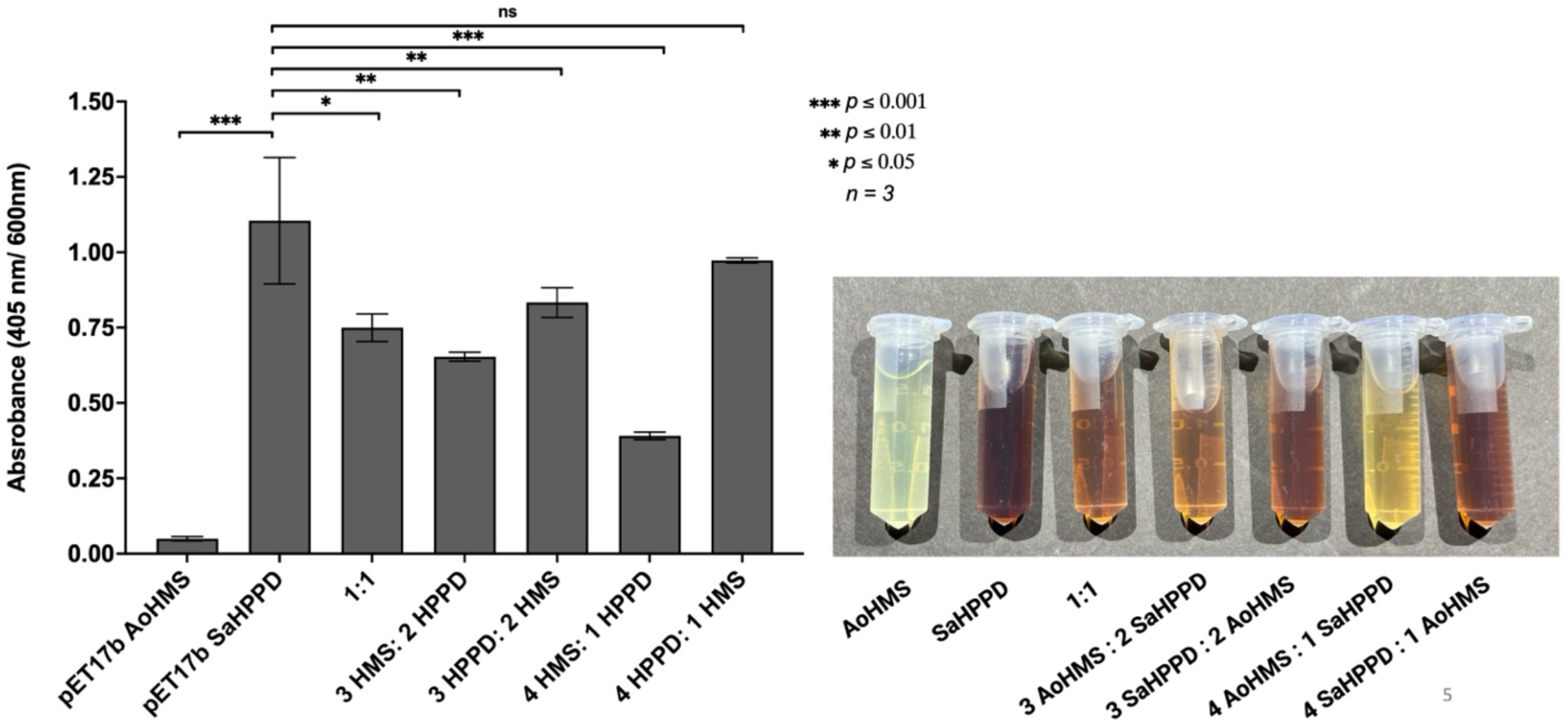
Creating a competition between *E. coli* strains harboring SaHPPD and AoHMS. Competition for tyrosine breakdown was develooped using two environmental microbial enzymes, AoHMS and SaHPPD in a mixed synthetic community of *E. coli* strains. The spent media from cultures of *E. coli* BL21(DE3) harboring AoHMS and SaHPPD in various ratios (1:1, 3:2, and 4:1) while induced with 1 mM IPTG and in the presence of 2 mM tyrosine showed pigment production. Quantification of pyomelanin production normalized by cell density for each sample is depicted in graphical format where ANOVA test was used to calculate statistical significance and p values, n=3. The p values for this graph indicate the significant or non-significant differences of pyomelanin produced in comparison the pigment production in the *E. coli* culture harboring SaHPPD.

To further understand this community dynamics, we carried out HPLC analysis to detect and measure secreted metabolites from the spent media. From the spent media of various cultures, HG, HMA and tyrosine were separated and measured. Figure 6 shows the quantification of expected metabolites from the individual and mixed culture of *E. coli* BL21(DE3) SaHPPD and *E. coli* BL21(DE3) AoHMS. When *E. coli* BL21(DE3) SaHPPD was grown separately, we found 1370 ± 440 µM of HG, which decreased gradually to 248 ± 10 µM as the ratio of SaHPPD to AoHMS changed from 4:1 to 1:4 in the mixed cultures (Table 1). On the other hand, the concentrations of HMA increased concurrently with decrease in HG. Additionally, the culture ratio of 1 SaHPPD:1 AoHMS, showed pigmentation that was between the pigmentation level of 3 SaHPPD:2 AoHMS and 2 SaHPPD:3 AoHMS, and we found equimolar concentration of HG and HMA in the spent media of this 1: 1 mixed culture. It is only possible to observe such results where ratios of metabolites are changing based on the changes in the abundance of enzymes, if the kinetic parameters of these enzymes are not vastly different which is the case here for HPPD and HMS^1,29–31,38,39^.

**Fig. 6:**
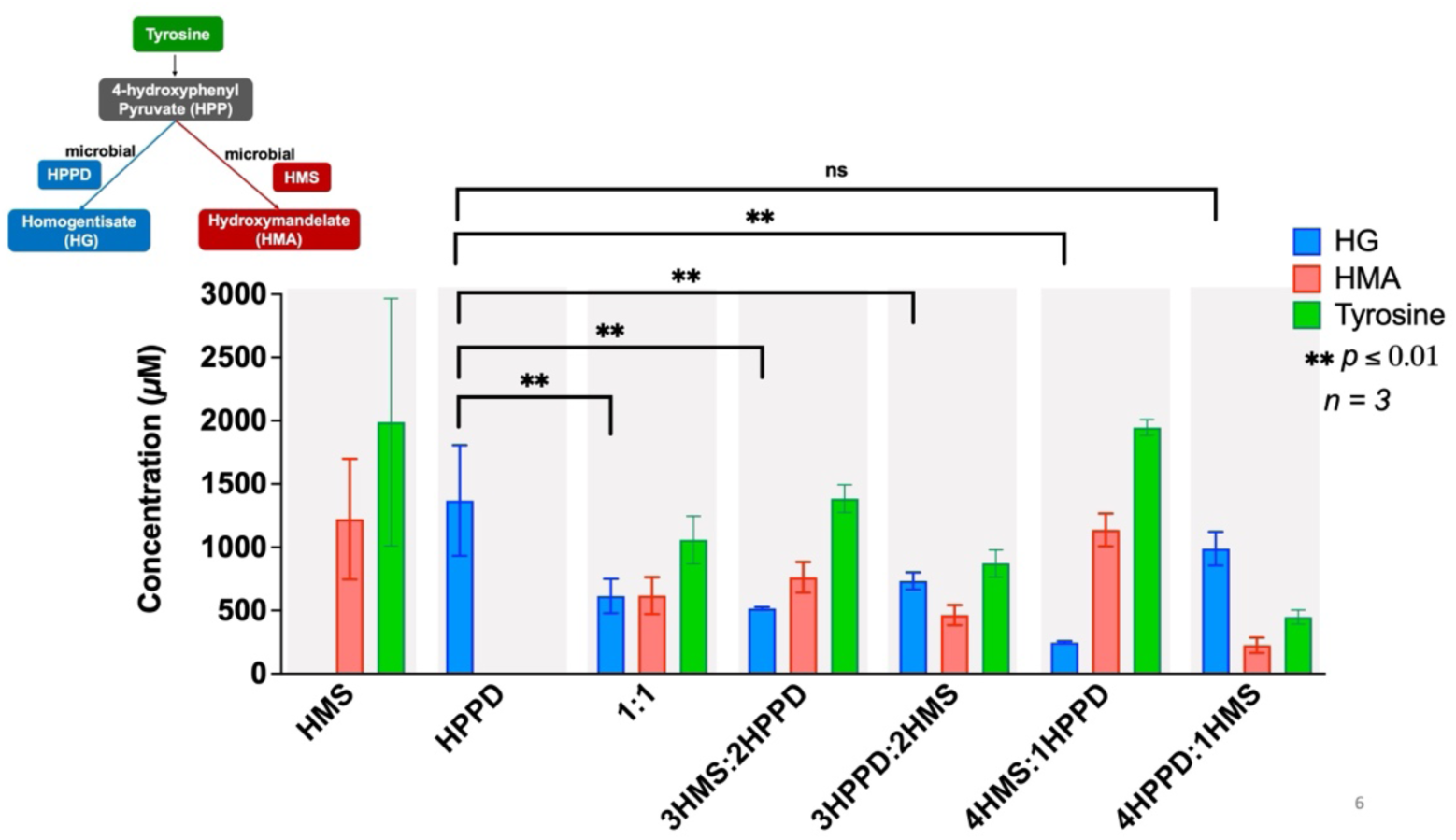
Quantification of metabolites from competition between *E. coli* strains harboring SaHPPD and AoHMS. The concentrations of secreted metabolites within the spent media of synthetic communities harboring SaHPPD and AoHMS at various ratios were measured. Each grey region represents an individual sample, while the different colors in each grey region represent separate metabolite. ANOVA test was used to calculate statistical significance and p values, n=3. The p values show the significance between HG production in samples where SaHPPD is in competition with AoHMS, compared to when SaHPPD is alone.

**Table 1.** Concentration of metabolites measured in individual and mixed cultures of *E. coli* BL21(DE3) SaHPPD and *E. coli* BL21(DE3) AoHMS.

| Culture | Homogentisate (HG) in $\mu\text{M}$ | Hydroxymandelate (HMA) in $\mu\text{M}$ | Tyrosine in $\mu\text{M}$ |
| --- | --- | --- | --- |
| HMS | N/A | 1220 ± 480 | 1989 ± 977 |
| HPPD | 1369 ± 437 | N/A | N/A |
| 4 HPPD:1 HMS | 988 ± 133 | 225 ± 29 | 1058 ± 188 |
| 3 HPPD:2 HMS | 732 ± 68 | 463 ± 27 | 1384 ± 110 |
| 2 HPPD:3 HMS | 516 ± 11 | 762 ± 56 | 872 ± 107 |
| 1 HPPD:4 HMS | 248 ± 10 | 1136 ± 124 | 1946 ± 64 |
| 1 HPPD:1 HMS | 614 ± 137 | 617 ± 49 | 448 ± 57 |

In contrast, if one of the enzymes was very fast, meaning it has a large Vmax, or had significantly high affinity for the substrate, meaning a low Km, then that would result in the predominance of the product of that enzyme even within the mixed cultures. In such a simple synthetic community, the competition for the resource, which is tyrosine, is thus driven directly by the competing enzyme activities. This shows that it is crucial to understand metabolic pathway diversions and kinetic parameters of enzymes involved in those pathways and such simplified synthetic microbial communities provides a unique strategy to predict metabolic outcomes. Interestingly, we observed large variations in the metabolite concentrations in pure cultures compared to mixed cultures (Fig. 6). Additionally, cultures with AoHMS showed a significant amount of tyrosine leftover or excess production in the spent media. Specifically in the case of AoHMS cultures, we saw around 2 mM tyrosine in the spent media which gradually decreased in mixed cultures with the correlating gradual increase in added HPPD. Whereas in pure SaHPPD cultures, we did not detect any tyrosine in the spent media. We think that there might be competition between AoHMS and an *E. coli* tyrosine aminotransferase enzyme (EcTAT) (Fig. 1). EcTAT is the only tyrosine catabolism enzyme present in *E. coli* cells which can catalyze a reversible conversion of tyrosine to 4-HPP and vice versa. If EcTAT is faster than AoHMS, then it might be catalyzing the conversion of 4-HPP to tyrosine faster than the conversion of 4-HPP to HMA by AoHMS and this might be resulting in the increased concentration of tyrosine in cultures that have AoHMS enzyme.

### Competition for tyrosine within a synthetic community of *E. coli* strains is driven by the activities of tyrosine metabolism enzymes

To further understand if the competitions in utilizing tyrosine are driven by enzyme activities and can such experiments be helpful in predicting metabolic outcomes with complex microbial communities, we created a synthetic community with *E. coli* BL21(DE3)pLysS harboring SaHPPD, *E. coli* BL21(DE3)pLysS harboring AoHMS, and *E. coli* Rosetta(DE3) harboring FjTAL (Fig. 7 and Fig. 8). *Flavobacterium johnsoniae* is an environmental microbe similar to *A. orientalis* and *S. avermitilis*^40,41^. *F. johnsoniae* produces yellow pigment by utilizing p-coumaric acid (pCA)^42^. pCA can be generated from tyrosine via the action of tyrosine ammonia lyase (TAL). In this community, tyrosine metabolic branching is occurring at two different points (Fig. 7 and Fig. 8). The first one is occurring at tyrosine, where FjTAL can pull it out of the homogentisate pathway and can directly convert it into p-coumaric acid (pCA) (Fig. 8). The second branching point is occurring at 4-HPP, where AoHMS can convert it into HMA (Fig. 8).

**Fig. 7:**
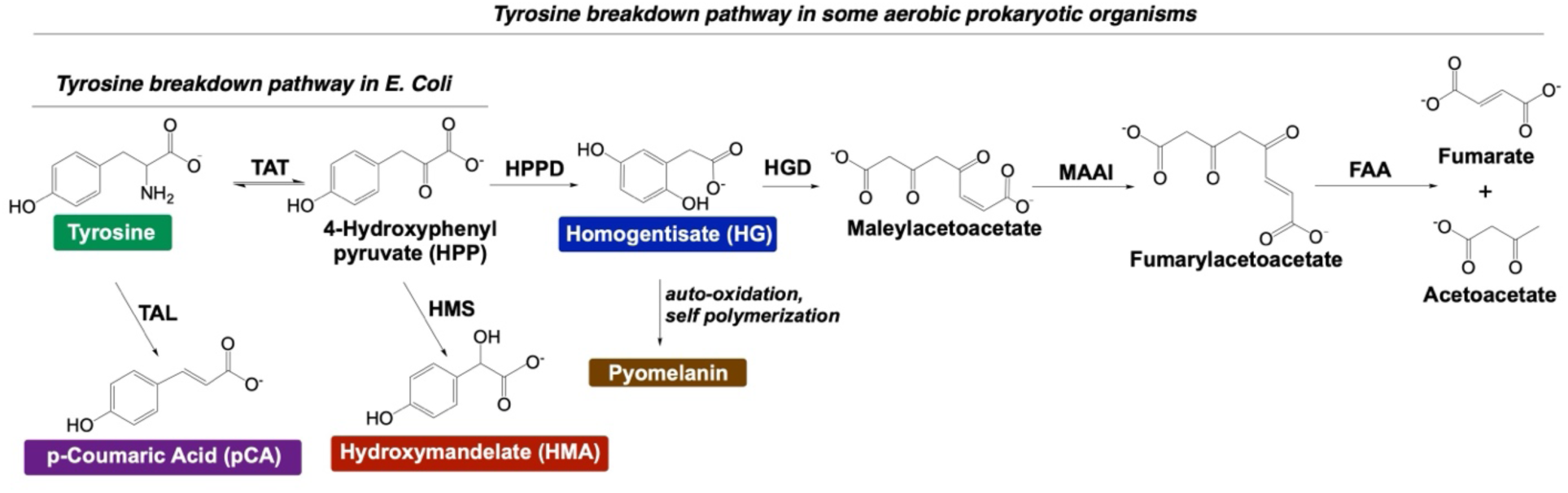
Tyrosine catabolism pathway diversion with two additional enzymes. The figure depicts the possible route for redirecting the tyrosine breakdown pathway by adding two additional enzymes involved in tyrosine metabolism from environmental microbes. Here, hydroxymandelate synthase (HMS) can convert tyrosine to hydroxymandelate (HMA) and tyrosine ammonia lyase (TAL) converts tyrosine to p-coumaric acid (pCA).

**Fig. 8:**
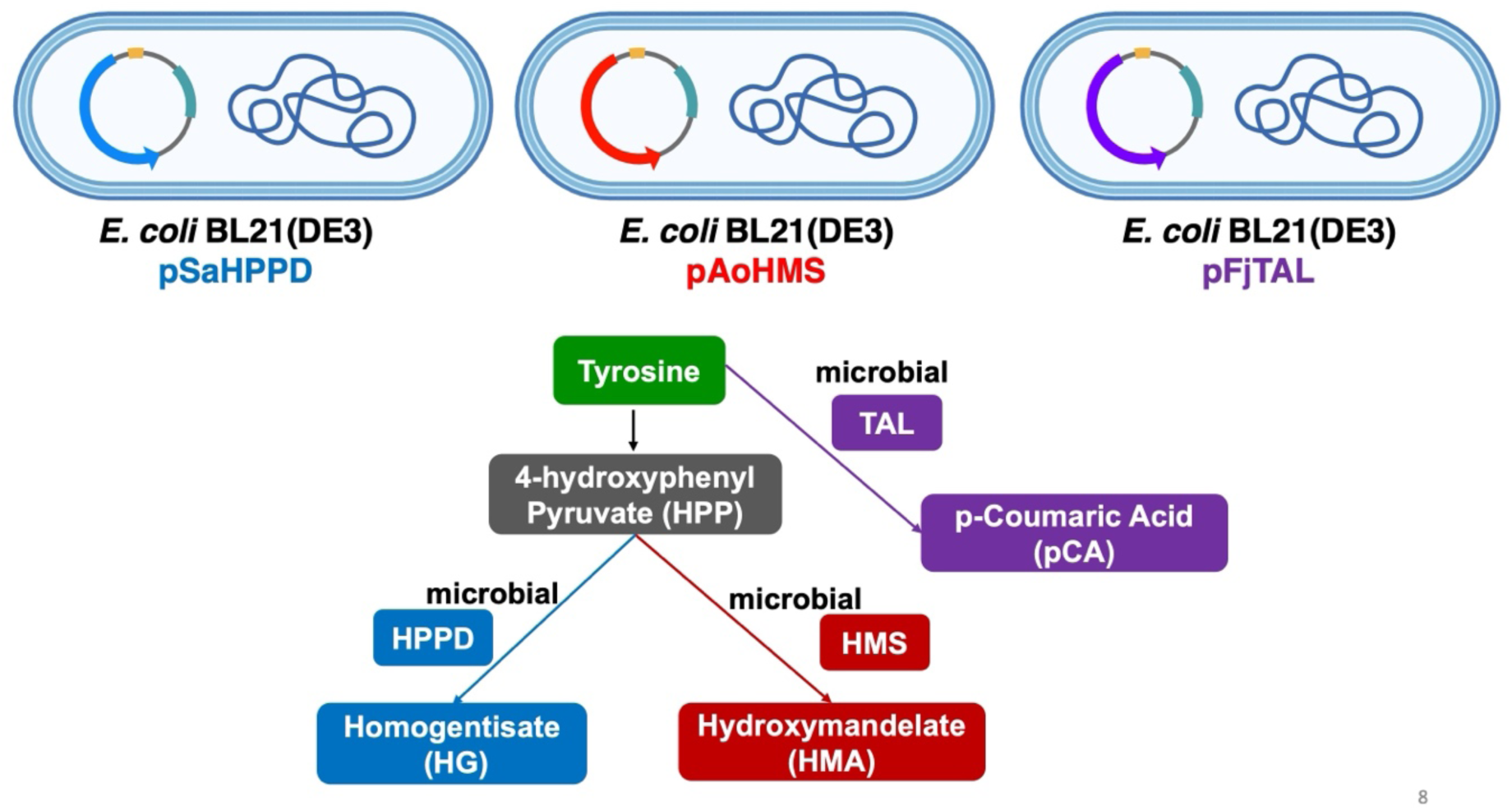
Generation of *E. coli* strains for Tyrosine catabolism pathway diversion with two additional enzymes. The figure depicts *E. coli* strains designed for redirecting the tyrosine breakdown pathway by adding two additional enzymes involved in tyrosine metabolism from environmental microbes. Here, hydroxymandelate synthase (HMS) can convert tyrosine to hydroxymandelate (HMA) and tyrosine ammonia lyase (TAL) converts tyrosine to p-coumaric acid (pCA).

Spent media of various mixed cultures and pure cultures were analyzed to detect and measure metabolites of tyrosine breakdown (Fig. 9). Here, while comparing the concentration of HG produced by SaHPPD to HG produced by the mixed culture of 1:1 ratio of SaHPPD:AoHMS, there was a significant decrease in HG in the mixed culture compared to the pure culture with only SaHPPD (Fig. 9). Moreover, we found that HG produced by SaHPPD was slightly higher than HMA produced by AoHMS in this 1:1 mix (Fig. 9), indicating higher activity coming from HPPD. We also observed around more than 1.5 mM of tyrosine in the spent media of 1:1 mix of AoHMS and SaHPPD. Such increase in tyrosine concentration in the mixed cultures of AoHMS and SaHPPD was observed before in our two-enzyme competing system experiments.

**Fig. 9:**
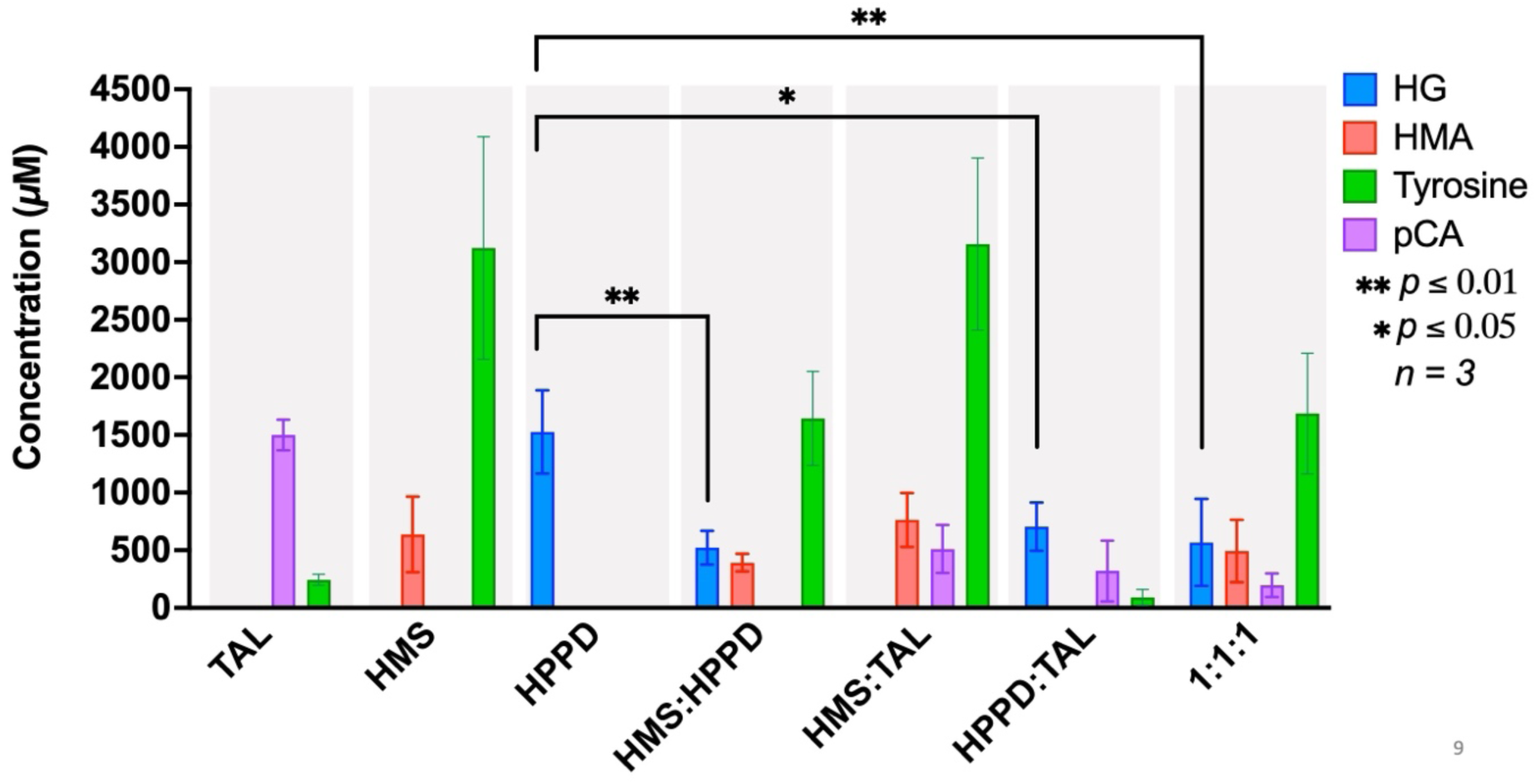
Quantification of metabolites from competition between *E. coli* strains harboring SaHPPD, AoHMS, and FjTAL. The concentrations of secreted metabolites within the spent media of synthetic communities harboring SaHPPD, AoHMS, and and fjTAL at various ratios were measured. Each grey region represents an individual sample, while the different colors in each grey region represent separate metabolite. ANOVA test was used to calculate statistical significance and p values, n=3. The p values show the significance between HG production in samples where SaHPPD is in competition with AoHMS and/or FjTAL, compared to the individual culture of SaHPPD.

Similarly, for the mixed culture of 1:1 ratio of SaHPPD:FjTAL, HG was decreased significantly in the mixed culture vs. pure culture. Analysis of the 1:1:1 mixture of SaHPPD:AoHMS:FjTAL showed the presence of almost equimolar concentrations of HG and HMA with two to three fold less pCA. In addition, the spent media showed the presence of almost 1.6 mM tyrosine. This mixture with the synthetic community of three different *E. coli* strains harboring three different enzymes, behaved very similarly to the 1:1 mix culture of HPPD:HMS. In this context, FjTAL did not provide much reduction of HG, especially while one of the goals was to decrease the amount of tyrosine going through the homogentisate pathway. Here we show that, AoHMS provided better metabolic diversion strategy if measured only by the change in HG levels. However, the huge increase seen in the tyrosine within the spent media of pure or mixed AoHMS cultures is not desirable. It will be interesting to understand if the increase in tyrosine is specific to *E. coli* cells specially in the presence of AoHMS.

### A competition between a human HPPD and gut microbial TAL can possibly provide a novel therapeutic strategy for metabolic disorders of tyrosine breakdown

Disorders that arise from tyrosine breakdown pathway severely impact the life of a patient even with the FDA approved therapy^21–24^. Although the medication given to these patients inhibits the tyrosine breakdown pathway, in the long run it can increase blood tyrosine concentrations which can lead to precipitation of tyrosine in the blood. For this reason, a therapeutic strategy where the tyrosine breakdown can be diverted from the mammalian homogentisate dependent pathway that stops the production of multiple harmful molecules under diseased conditions would provide a way to keep the blood tyrosine concentrations low (Fig. 10 and Fig. 11). To test this hypothesis, we expressed human HPPD enzyme (hHPPD), that catalyzes the conversion of 4-HPP to HG, and a tyrosine ammonia lyase (BoTAL) from the prominent gut microbe *Bacteroides ovatus,* that catalyzes the conversion of tyrosine to pCA, in *E. coli* BL21(DE3) cells (Fig. 11). Our pigmentation assay showed that the metabolic diversion was possible when tyrosine was partitioned between these two metabolic pathways (Fig. 12). We observed a notable decrease in the production of the pigment when *E. coli* BL21(DE3) strains containing hHPPD and BoTAL were mixed, which is a proxy for the decrease in the levels of secreted homogentisate (HG) (Fig. 2). This provided the first clue about the possibility of such metabolic diversions due to the competition between activities of a human enzyme and a gut microbial enzyme.

**Fig. 10:**
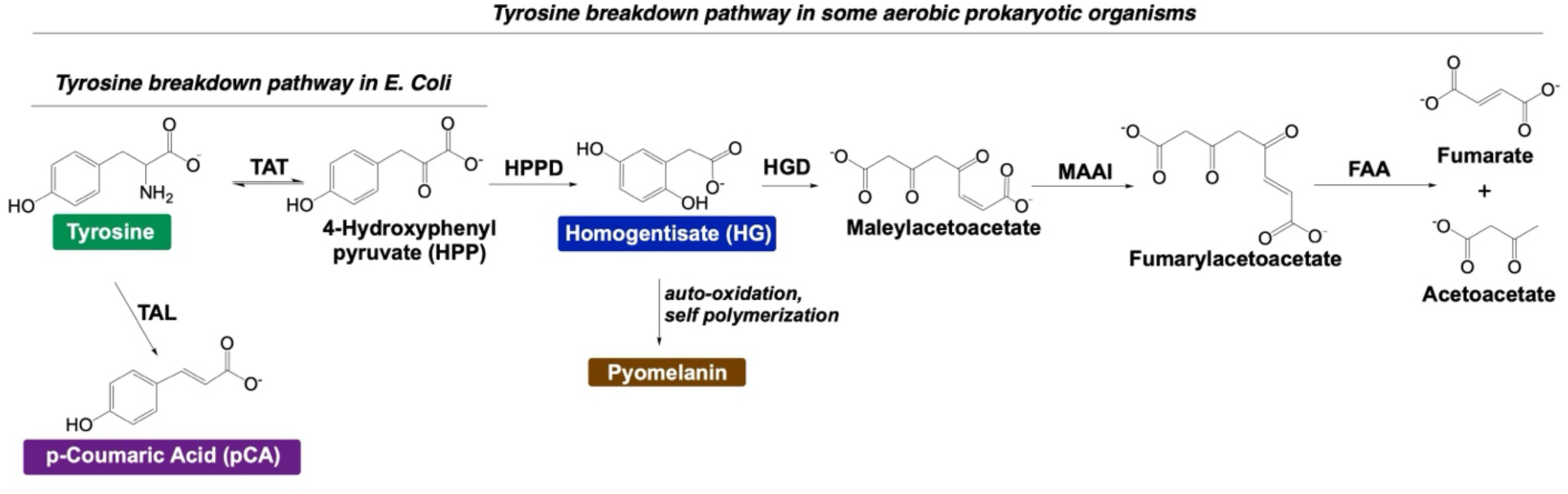
A possible mammalian tyrosine catabolism pathway diversion with a gut microbial enzyme. The figure depicts the possible route for redirecting the mammalian tyrosine breakdown pathway by adding a gut microbial enzyme involved in the tyrosine metabolism. Here, a gut microbial tyrosine ammonia lyase (TAL) converts tyrosine to p-coumaric acid (pCA).

**Fig. 11:**
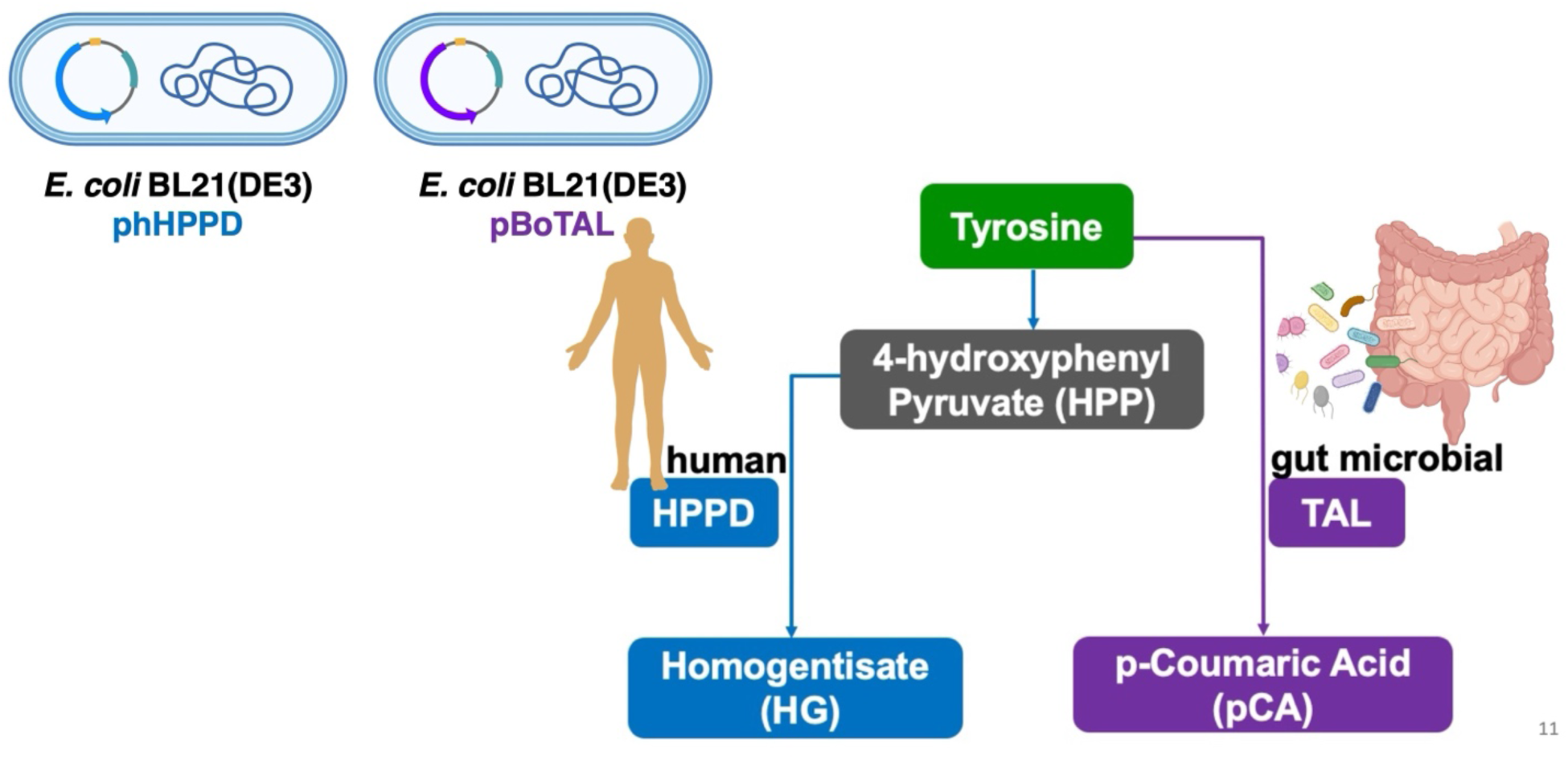
Generation of *E. coli* strains for mammalian tyrosine catabolism pathway diversion with a gut microbial enzyme. The figure depicts *E. coli* strains designed for redirecting the mammalian tyrosine breakdown pathway by adding a gut microbial enzyme involved in the tyrosine metabolism. Here, a gut microbial tyrosine ammonia lyase (TAL) converts tyrosine to p-coumaric acid (pCA).

**Fig. 12:**
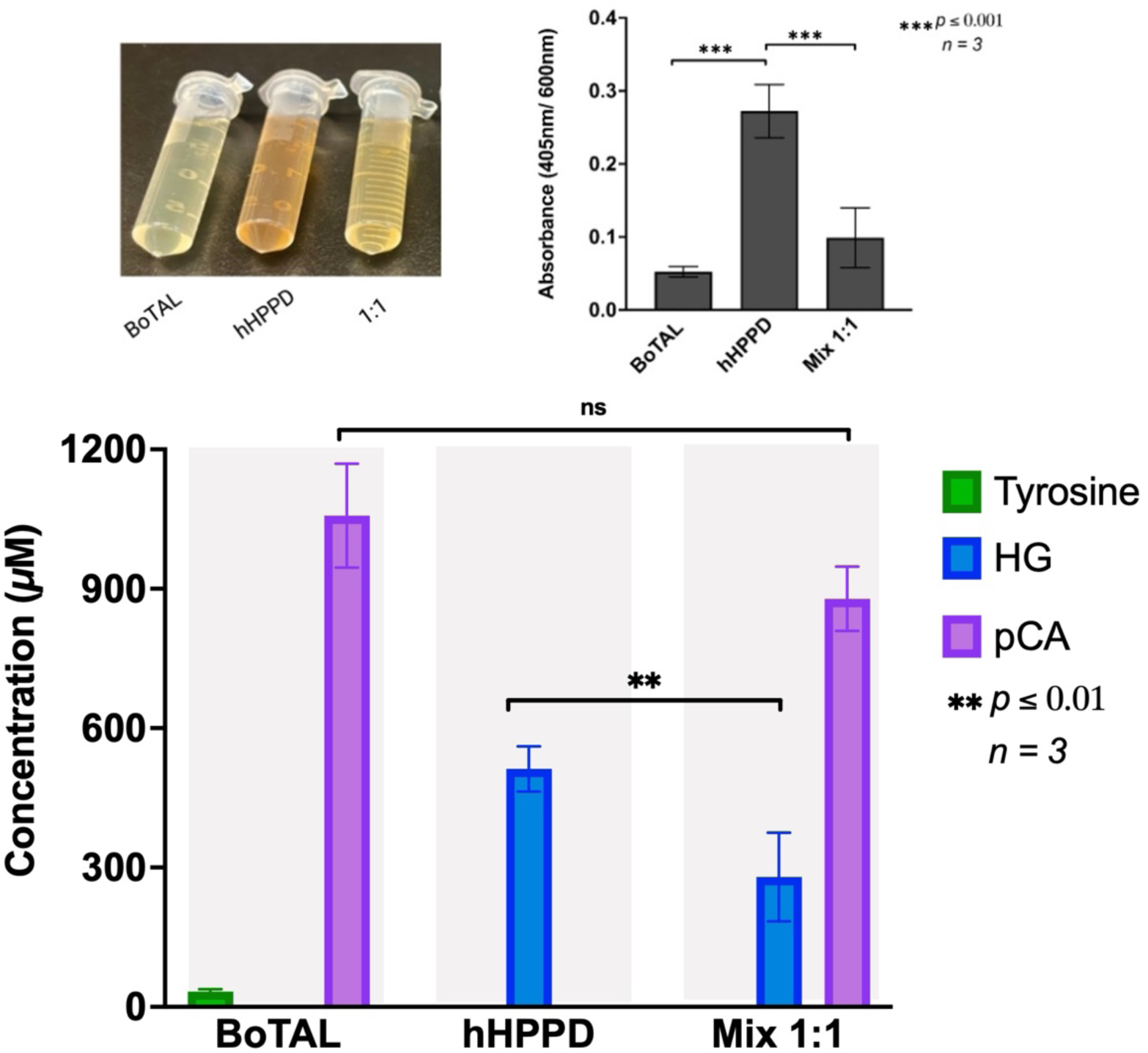
Creating a competition between *E. coli* strains harboring hHPPD and BoTAL. The spent media from cultures of *E. coli* BL21(DE3) harboring BoTAL and hHPPD while induced with 1 mM IPTG and in the presence of 2 mM tyrosine were utilized to test pigment production. Quantification of pyomelanin production normalized by cell density for each sample is presented in graphical format where ANOVA test was used to calculate statistical significance and p values, n=3. The p values for this graph indicate the significant or non-significant differences of pyomelanin produced in comparison the pigment production in the *E. coli* culture harboring hHPPD. Additionally, concentrations of secreted metabolites within the spent media of synthetic communities harboring hHPPD and BoTAL as compared to individual cultures were measured. Each grey region represents an individual sample, while the different colors in each grey region represent separate metabolite. ANOVA test was used to calculate statistical significance and p values, n=3. The p values show the significance between HG production in samples where hHPPD is in competition with BoTAL, compared to when hHPPD is alone.

Furthermore, the analysis of spent media provided additional information about the nature of this competition. We observed a marked decrease in the production of HG when cultures expressing hHPPD and BoTAL were mixed at 1:1 ratio (Fig. 12). Additionally, there was no tyrosine found in the spent media from the 1:1 mixed culture. These results show that, two basic requirements to create a potential therapeutic strategy were met here. The first where we detected a complete conversion of tyrosine during competition which provides a way to handle the insolubility of tyrosine. The second where due to the diversion from HG pathway to the pCA pathway, the concentrations of harmful metabolites produced via faulty tyrosine catabolism involving HG pathway would subsequently possibly be reduced.

## CONCLUSION

Through this study, we show that the delicate balance of tyrosine metabolites is controlled by the competitive interactions within different metabolic pathways, which can possibly encompass both inter-microbial competition and the dynamic between microbes and their host. The contest for tyrosine can potentially impact microbial community structures in various environments, while a similar competition in the human gut may have implications for human health. This study has highlighted the complexity of tyrosine metabolism within synthetic microbial communities, emphasizing the influence of specific enzymatic activities. Furthermore, we have demonstrated a unique metabolic redirection of tyrosine degradation by leveraging a competition between a human enzyme and a gut microbial enzyme, suggesting a promising approach for the development of therapeutics in future. This could possibly offer innovative treatments for metabolic disorders linked to tyrosine metabolism which will need to be experimentally tested.

## MATERIALS AND METHODS

### Materials

All restriction enzymes, competent cells and Monarch® Plasmid Miniprep Kit were purchased from New England BioLabs (NEB). Isopropyl β-D-1-thiogalactopyranoside (IPTG) and L-tyrosine disodium salt dihydrate 98% were both purchased from Sigma Aldrich. Costar 96 well, flat, clear-bottom plate were from fisher scientific. Precision Plus protein standards ladder, in a BIO-RAD Mini-PROTEAN TGX gel within a Mini-PROTEAN Tetra Cell holding 1X Tris/Glycine/Sodium dodecyl sulfate polyacrylamide (SDS) Buffer. Competent cells of *E. coli* BL21(DE3)pLysS and Rosetta(DE3) were from novagen. DL-4-hydroxymandelic acid monohydrate and homogentisic acid were from TCI Chemicals. p-coumaric acid and M9 minimal media were purchased from Sigma Aldrich.

### Experimental methods

#### Pyomelanin production in mixed cultures of *E. coli* BL21(DE3) with pET17b-SaHPPD and *E. coli* BL21(DE3) with pET17b-AoHMS

The genes for 4-hydroxyphenylpyruvate dioxygenase (SaHPPD) from *Streptomyces avermitilis* and hydroxymandelate synthase (AoHMS) from *Amycolatopsis orientalis* cloned in pET17b vector were received as a gift from Dr. Graham R. Moran. SaHPPD was cloned between NdeI and BamHI sites, whereas AoHMS was cloned between NdeI and XhoI sites. These plasmids were transformed into two separate *E. coli* strains, NEB 5⍺ and BL21(DE3). These cells were stored at - 80 °C as glycerol stocks.

After streaking the 20% glycerol stocks of both *E. coli* BL21(DE3) pET17b SaHPPD and *E. coli* BL21(DE3) pET17b AoHMS on individual LB agar with 100 µg/mL ampicillin plates, they incubated overnight in 37°C to have single colony formation. Single colonies of *E. coli* BL21(DE3) containing pET17b-SaHPPD and *E. coli* BL21(DE3) containing pET17b-AoHMS were used to start 5 mL starter cultures in LB with containing 100 µg/mL ampicillin. These cultures grew overnight at 37 °C with shaking at 200 rpm. Subsequently, these starter cultures were used to inoculate 50 mL cultures in LB containing 100 µg/mL ampicillin. The cultures were monitored by measuring OD600 for growth. Once the OD600 reached 0.6, these larger cultures were then subdivided into several 50 mL conical tubes containing 2 mM tyrosine and 1 mM IPTG in a total volume of 5 mL. 5 mL cultures of several different combinations of *E. coli* BL21(DE3) containing pET17b-SaHPPD and *E. coli* BL21(DE3) containing pET17b-AoHMS were created as follows: HMS : HPPD at 1:1, 3 HMS: 2 HPPD, 3 HPPD: 2 HMS, 4 HMS: 1HPPD, and 4 HPPD : 1 HMS. These test cultures were then placed at a slant on a shaker at 230 rpm and were cultivated at 37 °C for 24 h. After the 24 h of incubation, the cultures were analyzed by recording the OD600 (600 nm) and pigmentation (405 nm) absorbance values via microplate reader. For pigmentation measurements, cultures were centrifuged and then the supernatants underwent absorbance measurements at 405 nm. There were three independent replicates (n=3) for this experiment. Graphpad Prism was used to generate a bar graph from OD600 and 405 nm absorbance values, standard deviations and to conduct an ANOVA test to verify statistical significance. Pigmentation measurements at 405 nm were normalized to the cell density (OD) measured at 600 nm. This is to make sure that the observed pigmentation in various cultures is not due to the differences in the cell growth.

#### Cloning of AoHMS and SaHPPD into pET28a

SaHPPD and AoHMS were then subcloned in pET28a. SaHPPD was cloned between NdeI and BamHI sites, whereas AoHMS was cloned between NdeI and XhoI sites and transformed into *E. coli* NEB DH5⍺ strains. Plasmids containing both SaHPPD and AoHMS underwent DNA sequencing at plasmidsaurus. After sequence verification, both plasmids were transformed into *E. coli* BL21(DE3) strain for expression. Additionally, pET28a SaHPPD and pET28a AoHMS plasmids were transformed into another *E. coli* BL21(DE3) derivative, *E. coli* BL21(DE3)pLysS, where the competent cells were purchased from NEB.

#### Pyomelanin production from *E. coli* BL21(DE3) and *E. coli* BL21(DE3) pLysS strains harboring pSaHPPD

*E. coli* BL21(DE3) containing pET28a-SaHPPD and *E. coli* BL21(DE3)pLysS containing pET28a-SaHPPD were individually grown in 5 mL LB containing 50 µg/mL kanamycin. Cultures were grown within 50 mL conical tubes overnight at 37 °C at a slant with shaking at 230 rpm. These initial cultures were used to inoculate larger 50 mL subcultures in LB containing 50 µg/mL kanamycin within 125 mL Erlenmeyer flasks that grew at 37 °C with shaking at 230 rpm. Their growth was monitored by measuring culture density at OD600 using a Costar 96 well, flat, clear-bottom plate on a microplate reader. and a at 600 nm until they reached 0.6 optical density (OD). Once the OD600 was reached, 5 mL of the subcultures were aliquoted into four different 50 mL conical tubes per *E. coli* strain containing SaHPPD which were designed for different testing conditions. The four conditions were; (1) without IPTG or tyrosine, (2) with 1 mM IPTG, (3) with 2 mM tyrosine, and (4) with both 1 mM IPTG and 2 mM tyrosine. These cultures were incubated at 37 °C with shaking at 230 rpm for 24 hours. After the 24 h incubation, 2 mL of each culture form each condition was pipetted into sterile 2 mL eppendorfs and underwent centrifugation at 1000*g* for around ten minutes to collect supernatants. These supernatants were then transferred to a 96-well plate to analyze the brown pigment (pyomelanin) production within each test condition by measuring absorbance at 405 nm via a microplate reader. Separated pellets were stored in - 80°C until further use. There were three independent replicates (n=3) for this experiment. Graphpad Prism was used to generate a bar graph from OD600 and 405 nm absorbance values, standard deviations and to conduct an ANOVA test to verify statistical significance. Pigmentation measurements at 405 nm were normalized to the cell density (OD) measured at 600 nm. This is to make sure that the observed pigmentation in various cultures is not due to the differences in the cell growth.

#### Protein expression analysis via SDS-PAGE from *E. coli* BL21(DE3) and *E. coli* BL21(DE3)pLysS strains harboring pSaHPPD

The pellets collected from previous pigmentation cultures were thawed on ice and resuspended in 1.5 mL of sterile 1X M9 minimal media (sterile buffer). The optical densities of these resuspended samples were normalized and then 15 µL of each sample was mixed with 15 µL of Laemmli sample buffer containing dye and beta-mercaptoethanol (BME). The sample were placed in boiling water bath for ten minutes and then centrifuged for 5 min at 1000*g* and finally loaded alongside the Precision Plus protein standards ladder, in a BIO-RAD Mini-protean TGX gel for SDS-PAGE analysis. After the run the gel was stained with QC colloidal coomassie stain and was imaged with a BIO-RAD gel imager to confirm the protein band sizes.

#### Pyomelanin production in mixed cultures of *E. coli* BL21(DE3)pLysS containing pET28a-SaHPPD, *E. coli* BL21(DE3)pLysS pET28a-AoHMS, and *E. coli* Rosetta(DE3) containing pET28a-FjTAL

The gene for tyrosine ammonia lyase (FjTAL) (Gene ID: 644740509) from *Flavobacterium johnsoniae* was PCR amplified from the *F. johnsoniae* genome and cloned into pET28a vector between NdeI and XhoI restriction sites. This newly made pET28a-FjTAL was then transformed into *E. coli* Rosetta(DE3). pET28a-FjTAL was transformed to Rosetta to assure proper expression of FjTAL due to its ability harbor tRNAs for expression of proteins that are non-native to E. coli or not codon optimized. Additionally, due the requirement of chloramphenicol for *E. coli* Rosetta(DE3), we subcloned both SaHPPD and AoHMS in the *E. coli* BL21(DE3)pLysS strain that is also resistant to chloramphenicol. All three of these strains were plated on LB agar plates containing 50 µg/mL kanamycin and 25 µg/mL chloramphenicol to account for resistance against chloramphenicol and kanamycin resistance coming from pET28a vector. Plasmids from all three strains were sequenced through Plasmidsaurus, then 20 % glycerol stocks were created and stored in -80°C until further use.

Competition assays were carried out similar to AoHMS and SaHPPD competition assays as mentioned above. Single colonies form the each of the three strains mentioned above were used to create three 5 mL LB starter cultures containing 50 µg/mL kanamycin and 25 µg/mL chloramphenicol. These cultures grew overnight at 37 °C with shaking at 230 rpm. Subsequently, these starter cultures were used to inoculate 50 mL cultures in LB containing 50 µg/mL kanamycin and 25 µg/mL chloramphenicol. The cultures were monitored by measuring OD600 for growth. Once the OD600 reached 0.6, these larger cultures were then subdivided into several 50 mL conical tubes containing 2 mM tyrosine and 1 mM IPTG in a total volume of 5 mL. 5 mL cultures of several different combinations of *E. coli* BL21(DE3)pLysS containing pET28a-SaHPPD, *E. coli* BL21(DE3)pLysS containing pET28a-AoHMS, and *E. coli* Rosetta(DE3) containing pET28a-FjTAL were created as follows: only TAL, only HMS, only HPPD, HMS: HPPD at 1:1, TAL: HMS at 1:1, TAL:HPPD at 1:1, and HPPD:HMS:TAL at 1:1:1. These test cultures were then placed at a slant on a shaker at 230 rpm and were cultivated at 37 °C for 24 h. After the 24 h of incubation, the cultures were analyzed by recording the OD600 (600 nm) and pigmentation (405 nm) absorbance values via microplate reader. For pigmentation measurements, cultures were centrifuged and then the supernatants underwent absorbance measurements at 405 nm. There were three independent replicates (n=3) for this experiment. Graphpad Prism was used to generate a bar graph from OD600 and 405 nm absorbance values, standard deviations and to conduct an ANOVA test to verify statistical significance. Pigmentation measurements at 405 nm were normalized to the cell density (OD) measured at 600 nm. This is to make sure that the observed pigmentation in various cultures is not due to the differences in the cell growth.

#### Pyomelanin production in mixed cultures of *E. coli* BL21(DE3) containing pET28a-BoTAL and *E. coli* BL21(DE3) containing pET28a-hHPPD

Genes for tyrosine ammonia lyase (BoTAL) (Gene ID: 29455038) from *Bacteroides ovatus* (BoTAL) and human 4-hydroxyphenylpyruvate dioxygenase (hHPPD) (Gene ID: 3242), synthesized and codon optimized for *E. coli* expression, were received from GeneWiz. These genes were then cloned into the pET28a vector, where BoTAL was cloned between NdeI and BamHI sites and hHPPD was cloned between NcoI and BamHI. Both vectors pET28a-hHPPD and pET28a-BoTAL were then transformed into *E. coli* NEB DH5*⍺* and *E. coli* BL21(DE3) competent cells and glycerol stocks were stored in -80°C until further use.

The same procedures as previously stated above were used to analyze pyomelanin production with *E.coli* BL21(DE3) containing pET28a-hHPPD and *E.coli* BL21(DE3) containing pET28a-BoTAL mixed culture competition assays. The combinations for mixed cultures of *h*HPPD and *Bo*TAL were; only TAL, only HPPD, and HPPD:TAL at 1:1, 3 TAL: 2 HPPD, 3 HPPD: 2 TAL, 4 TAL: 1 HPPD, 4 HPPD: 1 TAL. These test cultures were then placed at a slant on a shaker at 230 rpm and were cultivated at 37 °C for 24 h. After the 24 h of incubation, the cultures were analyzed by recording the OD600 (600 nm) and pigmentation (405 nm) absorbance values via a microplate reader. For pigmentation measurements, cultures were centrifuged and then the supernatants underwent absorbance measurements at 405 nm. There were three independent replicates (n=3) for this experiment. Graphpad Prism was used to generate a bar graph from OD600 and 405 nm absorbance values, standard deviations and to conduct an ANOVA test to verify statistical significance. Pigmentation measurements at 405 nm were normalized to the cell density (OD) measured at 600 nm. This is to make sure that the observed pigmentation in various cultures is not due to the differences in the cell growth.

#### HPLC of secreted tyrosine metabolites from various *E. coli* strains in mixed competition assay cultures

For detection and measurement of secreted tyrosine metabolites, HPLC analysis of spent media from mixed cultures was carried out. Previously described competition assays of mixed cultures were carried out as mentioned above but instead of using LB media, the assays were conducted in M9 minimal media. Induced cultures containing 2 mM tyrosine were incubated for 48 h at 37 °C with shaking at 230 rpm. After 48 h of incubation, cells were harvested by centrifugation for all cultures and the spent media (supernatants) were collected. These supernatants (spent media) were then passed through 0.2 µm PES syringe filters to remove any cell debris and then through 10 kDa MWCO filters by centrifugation at 1000*g* to remove any small peptides and proteins. The flow through from these samples were then analyzed by HPLC as a final volume of 100 µl. Each sample contained 25 μM of Thiamine HCl as an internal standard.

Separation was carried out using a Synergi 4 μm Hydro-RP 80 Å 250 x 4.6 mm column on Waters e2695 separation module with a diode array detector via two separate methods to analyze peaks representing tyrosine, hydroxymandelic acid (HMA), homogentisate (HG), and p-coumaric acid (pCA). For isocratic method, samples were separated under isocratic conditions with a mobile phase consisting of 20 mM sodium phosphate, pH 2 and 0.5% acetonitrile at 1 mL/min^38^. For gradient method, mobile phase consisting of methanol-water with 0.1% formic acid (Buffer A, methanol with 0.1% formic acid and Buffer B, water with 0.1% formic acid). Briefly the method consisted of 5:95 (A:B) for 0-5 min, 5:95 to 20:80 from 5-14 min, 20:80 to 80:20 from 14-18 min, 80:20 from 18-25 min^43^. After 25 min, column was brought back to the initial conditions before loading the new sample. Samples were monitored using DAD and all metabolites could be visualized at 276 nm.

The peak areas from chromatograms of various samples were analyzed using the chromatography data software, Empower. Concentration of metabolites were calculated from peak areas by standard curves generated for tyrosine, HMA, HG, and pCA. Finally, the values were entered into GraphPad Prism to generate a plot showing the relationship between concentrations within different mixed samples. GraphPad Prism was utilized to generate bar graphs, to compute standard deviation, and to perform ANOVA testing to provide statistical analysis of data and provide p values.

## ACKNOWLEDGEMENTS

We would like to thank Dr. Graham R. Moran for providing vectors containing SaHPPD and AoHMS.

## AUTHOR CONTRIBUTIONS

DS and MM conceptualized the experiments. MM performed majority of the experiments. CP cloned a gene for FjTAL and performed sequence similarity analysis to find tyrosine ammonia lyases suitable for this study. MM prepared majority of the figures. DS edited figures. DS and MM wrote the manuscript. CP reviewed the manuscript and provided edits.

## DATA AVAILABILITY

All data of this study are included within the manuscript.

## REFERENCES

1 Shah, D. D. & Moran, G. R. in *2-Oxoglutarate-Dependent Oxygenases* (eds R.P. Hausinger & C.J. Schofield) Ch. 4-Hydroxyphenylpyruvate dioxygenase and hydroxymandelate synthase:2-oxo acid-dependent oxygenases of importance to agriculture and medicine., (RSC Publishing, 2015).

2 Dodd, D. et al. A gut bacterial pathway metabolizes aromatic amino acids into nine circulating metabolites. Nature 551, 648–652, doi:10.1038/nature24661 (2017).

3 Noda, S. & Kondo, A. Recent Advances in Microbial Production of Aromatic Chemicals and Derivatives. Trends Biotechnol 35, 785–796, doi:10.1016/j.tibtech.2017.05.006 (2017).

4 Shen, Y. P. et al. Recent Advances in Metabolically Engineered Microorganisms for the Production of Aromatic Chemicals Derived From Aromatic Amino Acids. Front Bioeng Biotechnol 8, 407, doi:10.3389/fbioe.2020.00407 (2020).

5 Summers, S. C. et al. The fecal microbiome and serum concentrations of indoxyl sulfate and p-cresol sulfate in cats with chronic kidney disease. J Vet Intern Med 33, 662–669, doi:10.1111/jvim.15389 (2019).

6 Harrison, M. A. et al. Identification of novel p-cresol inhibitors that reduce Clostridioides difficile’s ability to compete with species of the gut microbiome. Sci Rep 13, 9492, doi:10.1038/s41598-023-32656-8 (2023).

7 Machado Ribeiro, T. R., Brito, C. B. & Byndloss, M. X. Can our microbiome break our hearts? Collaborative production of p-cresol sulfate and indoxyl sulfate by commensal microbes increases susceptibility to thrombosis. mBio 15, e0269223, doi:10.1128/mbio.02692-23 (2024).

8 Schenck, C. A. & Maeda, H. A. Tyrosine biosynthesis, metabolism, and catabolism in plants. Phytochemistry 149, 82–102, doi:10.1016/j.phytochem.2018.02.003 (2018).

9 Schlune, A., Thimm, E., Herebian, D. & Spiekerkoetter, U. Single dose NTBC-treatment of hereditary tyrosinemia type I. J of Inher Metab Disea 35, 831–836, doi:10.1007/s10545-012-9450-9 (2012).

10 Thimm, E. et al. Neurocognitive outcome in patients with hypertyrosinemia type I after long-term treatment with NTBC. J of Inher Metab Disea 35, 263–268, doi:10.1007/s10545-011-9394-5 (2012).

11 Van Ginkel, W. G. et al. Long-Term Outcomes and Practical Considerations in the Pharmacological Management of Tyrosinemia Type 1. Pediatr Drugs 21, 413–426, doi:10.1007/s40272-019-00364-4 (2019).

12 Van Ginkel, W. G. et al. Blood and Brain Biochemistry and Behaviour in NTBC and Dietary Treated Tyrosinemia Type 1 Mice. Nutrients 11, 2486, doi:10.3390/nu11102486 (2019).

13 Phornphutkul, C. et al. Natural history of alkaptonuria. N Engl J Med 347, 2111–2121, doi:10.1056/NEJMoa021736 (2002).

14 Genovese, F. et al. Nitisinone Treatment Affects Biomarkers of Bone and Cartilage Remodelling in Alkaptonuria Patients. Int J Mol Sci 24, doi:10.3390/ijms241310996 (2023).

15 Wilcken, B. et al. Hawkinsinuria: a dominantly inherited defect of tyrosine metabolism with severe effects in infancy. N Engl J Med 305, 865–868, doi:10.1056/NEJM198110083051505 (1981).

16 Hocart, C. H., Halpern, B., Hick, L. A. & Wong, C. O. Hawkinsinuria--identification of quinolacetic acid and pyroglutamic acid during an acidotic phase. J Chromatogr 275, 237–243, doi:10.1016/s0378-4347(00)84371-6 (1983).

17 Tomoeda, K. et al. Mutations in the 4-hydroxyphenylpyruvic acid dioxygenase gene are responsible for tyrosinemia type III and hawkinsinuria. Mol Genet Metab 71, 506–510, doi:10.1006/mgme.2000.3085 (2000).

18 Brownlee, J. M., Heinz, B., Bates, J. & Moran, G. R. Product analysis and inhibition studies of a causative Asn to Ser variant of 4-hydroxyphenylpyruvate dioxygenase suggest a simple route to the treatment of Hawkinsinuria. Biochemistry 49, 7218–7226, doi:10.1021/bi1008112 (2010).

19 Conrad, J. A. & Moran, G. R. The interaction of hydroxymandelate synthase with the 4-hydroxyphenylpyruvate dioxygenase inhibitor: NTBC. Inorganica Chimica Acta 361, 1197–1201, doi:10.1016/j.ica.2007.07.036 (2008).

20 Lorquin, F., Piccerelle, P., Orneto, C., Robin, M. & Lorquin, J. New insights and advances on pyomelanin production: from microbial synthesis to applications. Journal of Industrial Microbiology and Biotechnology 49, kuac013, doi:10.1093/jimb/kuac013 (2022).

21 Ahmad, S., Teckman, J. H. & Lueder, G. T. Corneal opacities associated with NTBC treatment. Am J Ophthalmol 134, 266–268, doi:10.1016/s0002-9394(02)01532-5 (2002).

22 Schlune, A., Thimm, E., Herebian, D. & Spiekerkoetter, U. Single dose NTBC-treatment of hereditary tyrosinemia type I. J Inherit Metab Dis 35, 831–836, doi:10.1007/s10545-012-9450-9 (2012).

23 Thimm, E. et al. Neurocognitive outcome in patients with hypertyrosinemia type I after long-term treatment with NTBC. J Inherit Metab Dis 35, 263–268, doi:10.1007/s10545-011-9394-5 (2012).

24 Wisse, R. P., Wittebol-Post, D., Visser, G. & van der Lelij, A. Corneal depositions in tyrosinaemia type I during treatment with Nitisinone. BMJ Case Rep 2012, doi:10.1136/bcr-2012-006301 (2012).

25 Neuckermans, J., Mertens, A., De Win, D., Schwaneberg, U. & De Kock, J. A robust bacterial assay for high-throughput screening of human 4-hydroxyphenylpyruvate dioxygenase inhibitors. Sci Rep 9, 14145, doi:10.1038/s41598-019-50533-1 (2019).

26 Seo, D. & Choi, K.-Y. Heterologous production of pyomelanin biopolymer using 4-hydroxyphenylpyruvate dioxygenase isolated from Ralstonia pickettii in Escherichia coli. Biochemical Engineering Journal 157, 107548, doi:10.1016/j.bej.2020.107548 (2020).

27 Johnson-Winters, K., Purpero, V. M., Kavana, M., Nelson, T. & Moran, G. R. (4-Hydroxyphenyl)pyruvate dioxygenase from Streptomyces avermitilis: the basis for ordered substrate addition. Biochemistry 42, 2072–2080, doi:10.1021/bi026499m (2003).

28 Brownlee, J. M., Johnson-Winters, K., Harrison, D. H. & Moran, G. R. Structure of the ferrous form of (4-hydroxyphenyl)pyruvate dioxygenase from Streptomyces avermitilis in complex with the therapeutic herbicide, NTBC. Biochemistry 43, 6370–6377, doi:10.1021/bi049317s (2004).

29 Johnson-Winters, K., Purpero, V. M., Kavana, M. & Moran, G. R. Accumulation of multiple intermediates in the catalytic cycle of (4-hydroxyphenyl)pyruvate dioxygenase from Streptomyces avermitilis. Biochemistry 44, 7189–7199, doi:10.1021/bi047625k (2005).

30 Brownlee, J., He, P., Moran, G. R. & Harrison, D. H. Two roads diverged: the structure of hydroxymandelate synthase from Amycolatopsis orientalis in complex with 4-hydroxymandelate. Biochemistry 47, 2002–2013, doi:10.1021/bi701438r (2008).

31 He, P., Conrad, J. A. & Moran, G. R. The rate-limiting catalytic steps of hydroxymandelate synthase from Amycolatopsis orientalis. Biochemistry 49, 1998–2007, doi:10.1021/bi901674q (2010).

32 Chen, T. S. & Inamine, E. S. Studies on the biosynthesis of avermectins. Arch Biochem Biophys 270, 521–525, doi:10.1016/0003-9861(89)90534-1 (1989).

33 Chen, T. S., Inamine, E. S., Hensens, O. D., Zink, D. & Ostlind, D. A. Directed biosynthesis of avermectins. Arch Biochem Biophys 269, 544–547, doi:10.1016/0003-9861(89)90138-0 (1989).

34 Schulman, M. D., Valentino, D. & Hensens, O. Biosynthesis of the avermectins by Streptomyces avermitilis. Incorporation of labeled precursors. J Antibiot (Tokyo) 39, 541–549, doi:10.7164/antibiotics.39.541 (1986).

35 Rubinstein, E. & Keynan, Y. Vancomycin revisited - 60 years later. Front Public Health 2, 217, doi:10.3389/fpubh.2014.00217 (2014).

36 Hubbard, B. K., Thomas, M. G. & Walsh, C. T. Biosynthesis of L-p-hydroxyphenylglycine, a non-proteinogenic amino acid constituent of peptide antibiotics. Chem Biol 7, 931–942, doi:10.1016/s1074-5521(00)00043-0 (2000).

37 Omura, S. et al. Genome sequence of an industrial microorganism Streptomyces avermitilis: deducing the ability of producing secondary metabolites. Proc Natl Acad Sci U S A 98, 12215–12220, doi:10.1073/pnas.211433198 (2001).

38 Shah, D. D., Conrad, J. A., Heinz, B., Brownlee, J. M. & Moran, G. R. Evidence for the mechanism of hydroxylation by 4-hydroxyphenylpyruvate dioxygenase and hydroxymandelate synthase from intermediate partitioning in active site variants. Biochemistry 50, 7694–7704, doi:10.1021/bi2009344 (2011).

39 Shah, D. D., Conrad, J. A. & Moran, G. R. Intermediate partitioning kinetic isotope effects for the NIH shift of 4-hydroxyphenylpyruvate dioxygenase and the hydroxylation reaction of hydroxymandelate synthase reveal mechanistic complexity. Biochemistry 52, 6097–6107, doi:10.1021/bi400534q (2013).

40 Kim, J. J. et al. Flavobacterium compostarboris sp. nov., isolated from leaf-and-branch compost, and emended descriptions of Flavobacterium hercynium, Flavobacterium resistens and Flavobacterium johnsoniae. Int J Syst Evol Microbiol 62, 2018–2024, doi:10.1099/ijs.0.032920-0 (2012).

41 Chen, W. M., Huang, W. C., Young, C. C. & Sheu, S. Y. Flavobacterium tilapiae sp. nov., isolated from a freshwater pond, and emended descriptions of Flavobacterium defluvii and Flavobacterium johnsoniae. Int J Syst Evol Microbiol 63, 827–834, doi:10.1099/ijs.0.041178-0 (2013).

42 Schoner, T. A., Fuchs, S. W., Schonau, C. & Bode, H. B. Initiation of the flexirubin biosynthesis in Chitinophaga pinensis. Microb Biotechnol 7, 232–241, doi:10.1111/1751-7915.12110 (2014).

43 Cui, P. et al. Characterization of two new aromatic amino acid lyases from actinomycetes for highly efficient production of p-coumaric acid. Bioprocess Biosyst Eng 43, 1287–1298, doi:10.1007/s00449-020-02325-5 (2020).

